# Reconstructing the ecology of a Jurassic pseudoplanktonic megaraft colony

**DOI:** 10.1101/566844

**Authors:** Aaron W. Hunter, David Casenove, Emily G. Mitchell, Celia Mayers

## Abstract

Pseudoplanktonic crinoid megaraft colonies are an enigma of the Jurassic. They are among the largest in-situ invertebrate accumulations ever to exist in the Phanerozoic fossil record. These megaraft colonies and are thought to have developed as floating filter-feeding communities due to an exceptionally rich relatively predator free oceanic niche, high in the water column enabling them to reach high densities on these log rafts. However, this pseudoplanktonic hypothesis has never actually been quantitatively tested and some researchers have cast doubt that this mode of life was even possible. The ecological structure of the crinoid colony is resolved using spatial point process techniques and its longevity using moisture diffusion models. Using spatial analysis we found that the crinoids would have trailed preferentially positioned at the back of migrating structures in the regions of least resistance, consistent with a floating, not benthic ecology. Additionally, we found using a series of moisture diffusion models at different log densities and sizes that ecosystem collapse did not take place solely due to colonies becoming overladen as previously assumed. We have found that these crinoid colonies studied could have existed for greater than 10 years, even up to 20 years exceeding the life expectancy of modern documented megaraft systems with implications for the role of modern raft communities in the biotic colonisation of oceanic islands and intercontinental dispersal of marine and terrestrial species.

**Significance statement:** Transoceanic rafting is the principle mechanism for the biotic colonisation of oceanic island ecosystems. However, no historic records exist of how long such biotic systems lasted. Here, we use a deep-time example from the Early Jurassic to test the viability of these pseudoplanktonic systems, resolving for the first time whether these systems were truly free floating planktonic and viable for long enough to allow its inhabitants to grow to maturity. Using spatial methods we show that these colonies have a comparable structure to modern marine pesudoplankton on maritime structures, whilst the application of methods normally used in commercial logging is used to demonstrate the viability of the system which was capable of lasting up to 20 years.

## Introduction

Transoceanic rafting is a fundamental feature of marine evolutionary biogeography and ecology, often invoked to explain the origins of modern global patterns of species distributions (1, 2, 3, 4). These communities have been recorded today lasting up to 6 years (5). However, the deep time ecology of these communities has never been investigated in detail (6). In recent communities, such rafts have included highly adapted bivalves, barnacle, limpets, bryozoans, sea anemones, amphipods, and isopods (5). In the Jurassic these communities also consisted of specially adapted crinoids, whose apparent maturity suggests that these communities had to have lasted longer than modern examples (>6 years) (6). The structure and duration and these colonies has remained a mystery, with most studies choosing to focus on how the crinoids were adapted rather than the viability of the system, prompting intense debate on their lifestyle (7, 8) rather than the ecological structure and longevity of the habitat. This study uses the latest ecological techniques used in paleobiology to reconstruct the ecology and duration of these rafts.

Crinoids or sea lilies were a major part of the Jurassic shallow sea ecosystem, with crinoids found in a diverse suite of shallow marine environments (9). The monospecific crinoid colonies preserved on wood rafts are one of the most enigmatic and iconic of these communities (10). Found globally, they represent one of the largest *in-situ* invertebrate accumulations found in the fossil record (6), the only fossil example of transoceanic rafting with up to 100 individuals covering oyster-encrusted logs up to 14m long (11). An ongoing debate prevails on whether these crinoids could have colonized and persisted on these floating log habitats or they were instead part of benthic islands systems typical of the Mesozoic (12). Previous studies have not quantitatively addressed this quandary. In the present study we use spatial statistics and diffusion modelling in a novel approach to test whether this pseudoplanktonic mode of life existed. Spatial statistics are used to test whether the logs were colonized in open water or on the substrate, and diffusion models then quantify floating-log mechanics to test how long a floating system could have existed.

The spatial positions of *Seirocrinus* on one of the largest and best-preserved Early Jurassic floating wood examples known, the giant ‘Hauff Specimen’ from Holzmaden, Germany (13) were mapped (Fig. 1 and *SI Appendix*, Figs. S1–S3). The spatial patterns of benthic organisms depend on the dispersal of larvae (14, 15), the environmental conditions in which they settled, and whether the conditions were favorable for them to grow to adulthood (16, 17). Therefore, the spatial patterns of the crinoids can be used to try to “reverse engineer” the conditions in which they settled in order to deduce the environmental conditions (what part of the water column the log was in) when colonized (18, 19).

**Fig. 1.**
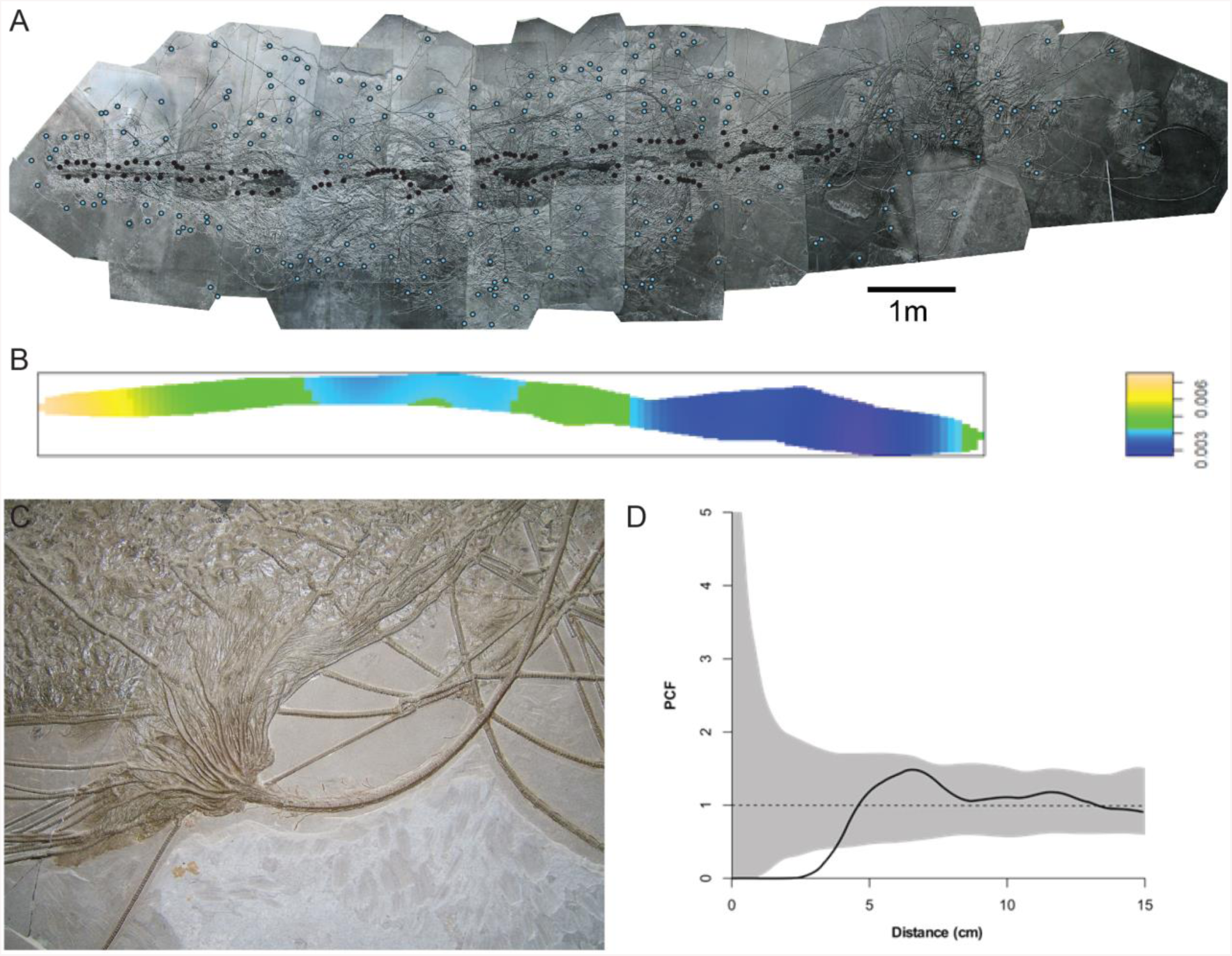
Crinoid fossil megaraft, the ‘Hauff Specimen’ from Holzmaden (G1). (*A*) Log with spatial analysis data points; key: blue= crinoid crowns, black= attachment discs. (*B*) Spatial analysis plot. (*C*) Close up view of crinoid crown and stem sections. (*D*) PCF distance plot.

This analysis was complemented by analyses using density/diffusion models (20, 21) to constrain the length of time logs of differing sizes and densities, with a full colony, could stay afloat as water infiltrates the wood over time. This diffusion analysis used three sets of size defined colonies (*SI Appendix*, Figs. S4–S9)), including small colony specimens (S1 and S2) (*SI Appendix*, Figs. S4 and S5) medium colonies (M1 and M2) (*SI Appendix*, Figs. S6 and S7) and finally massive giant colonies from Holzmaden (G1 “The Hauff Specimen” and G2) (*SI Appendix*, Figs. S8 and S9).

## Results

The spatial analyses clearly show an ecological signal of directionality along the log with the highest density on the left hand side, with decreasing density along the log (Fig. 1*B* and *SI Appendix*, Figs. S2 and S3). This directionality can be modelled by heterogeneous Poisson model depending on the *x* co-ordinate (*SI Appendix*, Table S1). The diffusion analysis is represented by two models, which constantly show that the largest of the log systems could have survived for a minimum of 2 years and a maximum of 20 years. (Fig. 2, and *SI Appendix*, Table S1) Thus allowing the crinoid colony to grow to maturity and confirming the viability of the pseudoplanktonic hypothesis. Firstly, the diffusion model assumes that the system is not adequately sealed. When the diffusion model is applied to the small logs (S1 and S2) (*SI Appendix*, Figs. S4 and S5) without the population, the log would sink after 800 days, however when the colony is added it would sink within 1 day (*SI Appendix*, Fig. S5). The apparent longevity of the logs is increased by their size, with the medium sized logs (M1 and M2) (*SI Appendix*, Figs. S6 and S7) surviving 100 days (800 without the colony) and the massive colonies (G1 and G2) (*SI Appendix*, Figs. S8 and S9) surviving up to 400 days (800 days without the colony) (*SI Appendix*, Fig. S9). Our results suggest that without any natural sealing the largest megarafts could only have survived for just over two years with the smaller systems unviable despite being relatively common. We propose that in order for the system to survive for long enough for the animals to grow to maturity the natural sealing of the wood structure would be needed with the oysters and the crinoids being part of efficient sealing of the system. The population model incorporated both the spatial distribution along the log as well as life-history estimates (fecundity, mortality, settling rates and maturation time) for a complete life cycle of extant oyster *Ostrea chilensis*. Simulations for the growth of oyster communities including spawning rate, per capita reproductive rate and mortality rate correspond to average values taken from the literature about *Ostrea chilensis*. The model allowed reconstruction of the number of years required to reach sinking density for the whole log for various values of K (*SI Appendix*, Tables S2 and S3). In some cases, the dimensions of the log combined with the density of the wood material would allow the oyster and crinoid loaded log never to reach sinking threshold within the 20-year window (Fig. 2).

**Fig. 2.**
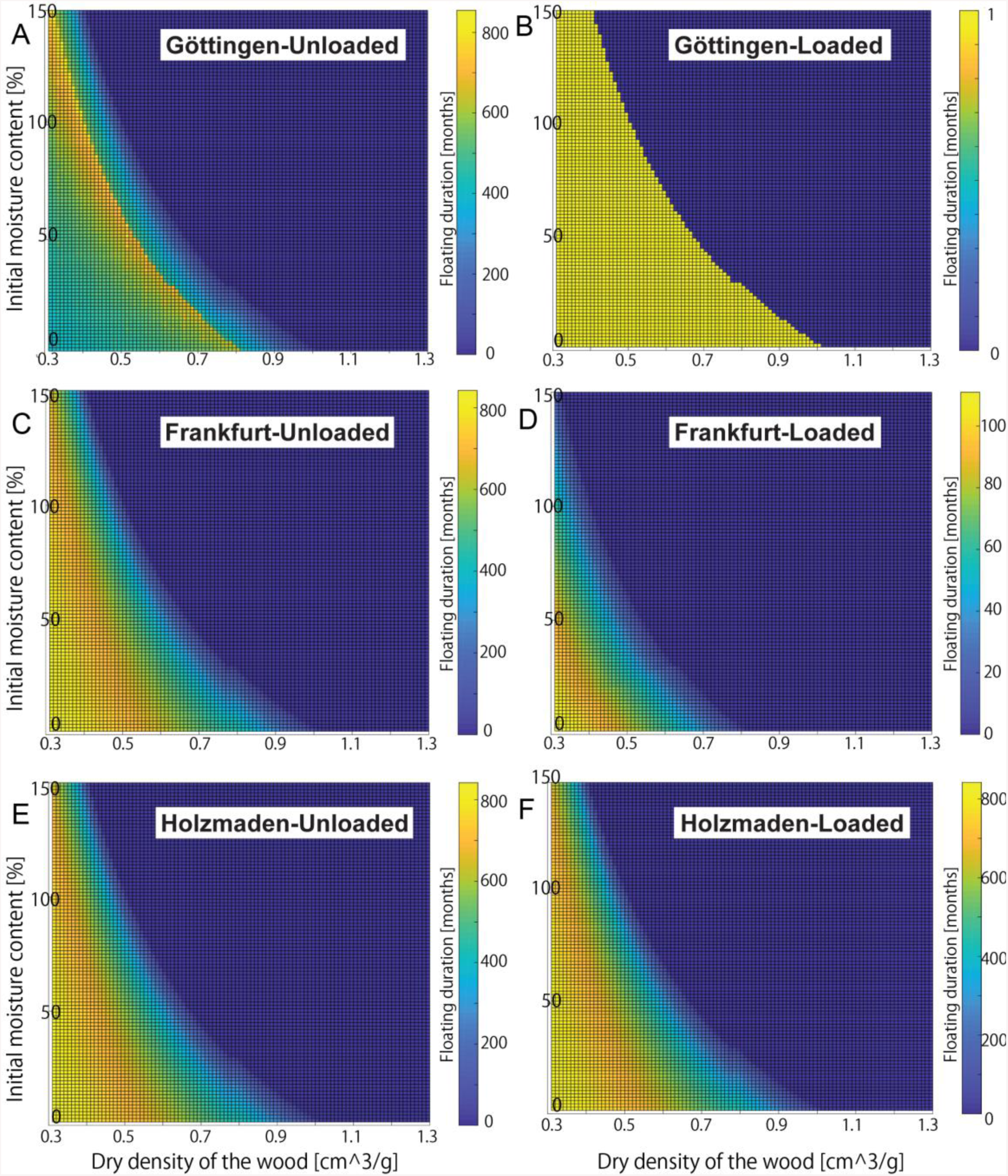
Wood diffusion gradient plots. (*A*) S1 unloaded. (*B*) S1 loaded. (*C*) M1 unloaded. (*D*) M1 loaded. (*E*) G1 unloaded. (F) G1 loaded.

## Discussion

The crinoids attached preferentially to one end of the log structure (Fig. 1*B*, and *SI Appendix*, Figs. S2 and S3), with other parts of the log more sparsely populated and attachment discs barely developed on the upper surface (Fig. 3).

**Fig. 3.**
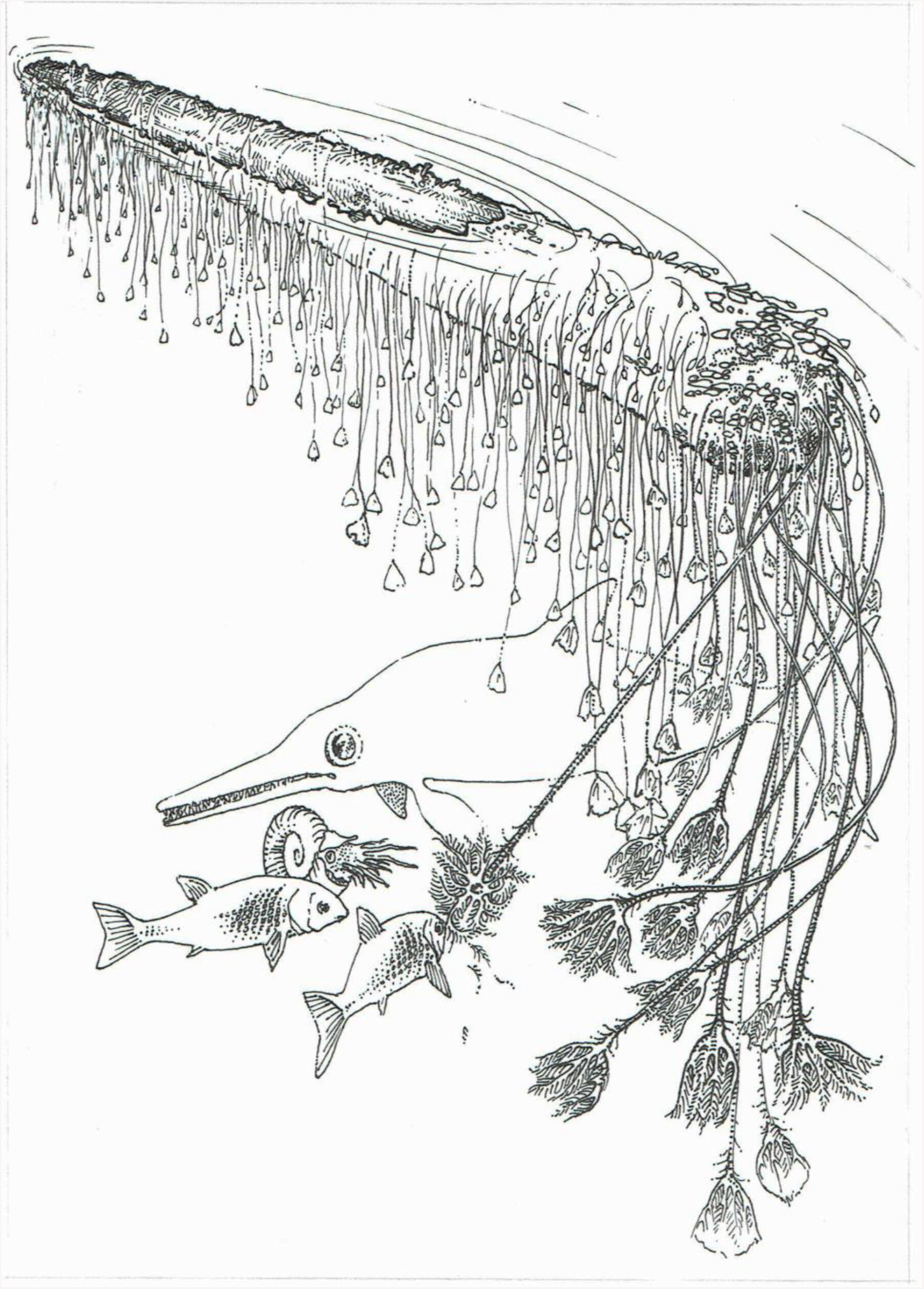
Reconstruction of the Holzmaden ‘Hauff Specimen’ crinoid megaraft colony (G1).

These spatial analyses support the hypothesis that our largest colony was structured as a floating body not a benthic structure. This feature is demonstrated by modern shipworms, for example, which preferentially accumulate at one end of the structure (22). In the case of a ship, this would be the back or stern. If the log were moving through the water even at a low rate the bow of the structure would still be subject to the most current pressure, so the most hospitable part of the structure would be the area of least resistance, the stern. However, this asymmetric distribution, combined with the observation that the vast majority of crinoids are attached to the underside of the log (Figs. 1*B* and 3 and *SI Appendix*, Figs. S2 and S3), would make the log unbalanced towards one end, bringing the colony’s stability into question. The conventional view is that the community would have rapidly sunk and been preserved at anoxic depths due to becoming unstable or overladen (13). However, our diffusion model does not support rapid sinking for three key reasons.

Firstly, the growing community of oysters and crinoids even at the climax state would still be a minor component of the total weight of the system. This total maximum weight (in very rare cases) would have varied from 630 to 880 kg depending on carry capacity and density compared to estimates of between 1100 kg to 15,000 kg for the total mas of the log depending on wood type (*SI Appendix*). Secondly, without efficient sealing our catastrophic models suggest the log could have supported the population of crinoids for a substantial amount of time. Even with a system designed to be a catastrophic scenario taking on water at a constant rate until the log raft system failed, the largest examples (G1 and G2) (*SI Appendix*, Figs. S8 and S9) could have survived for up to two years (*SI Appendix*, Table S3). Preserved fossil examples in this study were close to their maximum point viability as a mobile substrate. This point is evident from smaller logs (S1 and S2) (*SI Appendix*, Figs. S4 and S5)whose population was simply too big to be supported by the smaller log structure. However, the model looks at the viability of the climax community that would take time to develop, and would expand the life of the colony within the parameters of the model >2 years. However, modern growth rates for recent isocrinid crinoids 30-40 cm (23) in length suggests it would have taken at least 10 years for *Seirocrinus* to reach maturity. Although the growth rates of this crinoid may have been faster due to the unusual structure of the stem (24). For these communities to exist to maturity the log needs to float for much longer than 2 years (*SI Appendix*, Figs. S8 and S9). Our oyster community growth model in contrast, predicts that the megaraft was viable for up to and exceeding 20 years (Fig 2 and *SI Appendix*, Table S3). This length of time would require the structure to be sealed by an oyster coating to preserve any failures in the wood structure. The key to understanding how this lifespan of the megaraft was possible, is firstly by identifying areas of weaknesses were the structure is most likely to fail, this would include areas on the surface not coated in bark and secondly the ends of the log system that expose the wood xylem (25). However, it is clear from all the examples examined that most of the surface is covered in oysters (Fig. 1) and as our results suggest that the crinoids themselves are preferentially distributed at one terminal end of the system (Figs. 1*B* and 3 and *SI Appendix*, Figs. S2 and S3), contributing to the natural sealing of that end of the log system. Not all areas of the log were sealed and this would have shortened the lifespan of the community.

Thirdly, the viably of the community would have been influenced by the natural properties of the wood itself. Although it is hard to model the wood structure with no fossil remains available in the Posidonia Shale (26), the gymnosperm wood structure with the addition of the oyster coating and other agents such as aquatic fungus and algal slime must have allowed for sealing of the log and increased its longevity substantially beyond our models of continuous infiltration (27). Estimates vary for how long modern gymnosperm pine driftwood can survive (see *SI Appendix* for full discussion). Some estimates are as little as 17 months (28) while others suggest that the wood could have remained buoyant up to 5 years after entering the marine environment (see *SI Appendix*). The exceptional properties of the Jurassic gymnosperm wood with natural sealing by the oysters and crinoids, and without modern aggressive wood boring predators, which evolved later in the Mesozoic (29) the wood would have likely floated for longer than the 5 years observed in today’s driftwood. The colony could therefore have survived for the 10 years needed for a community to reach maturity, and possibly longer, until the decay and water infiltration of the wood reached a limit that could no longer support any further community growth. At which point the log sank and the crinoids died, being perfectly preserved by the anoxic conditions on the seabed (11) (Fig. 1*A* and 1*C*). Evidence suggests that fragments might have broken off while the system was in a state of decay which might explain the occurrence of single mature individuals attached to relatively small fragments of wood observed across the museum collections examined.

Pseudoplanktonic organisms are highly significant for global ecology and biodiversity, as a mechanism for global colonization (30), typically attaching themselves to necktic and planktonic organisms or other floating objects such as driftwood or flotsam. These are significant in today’s ecosystems and were responsible for the colonisation of oceanic islands such as Hawaii (31). These temporary rafts can become a permanent home to self-contained long-term communities such as, in modern ecosystems, external goose barnacles, tunicates and bryozoans (32), with wood boring bivalves such as shipworms inhabiting the internal structures (33).

Our analysis suggests that the exceptionally preserved Jurassic crinoid megarafts are the longest surviving communities to exist in the fossil record. Our results not only demonstrate that they have an ecological structure consistent with a recent living megaraft colony (22, 34, 35, 36) (see *SI Appendix*) but with efficient sealing these communities could have survived for up to and exceeding 20 years. In contrast, those recorded today have survived for up to 6 years. These extant megarafts of marine debris deliver substantial communities of adult organisms capable of reproduction or colonization by zooplankton in marginal marine environments from one continental margin to another (5). Rafts tend to be one-way arrival and deposition events which limit their journey time and life span before they encounter another system. These colonies are slow moving (1 to 2 knots) compared with modern commercial vessels (20 to >25 knots). These data are consistent with palaeoenvionmental interpretations suggesting an inhospitable seabed (37), indicating that these colonies remained afloat to survive and developed largely in isolation and could easily have spread around the globe. Our data suggests that the life of the colony was far more dependent on the wood structure than previously thought. Although the total weight of the crinoids would finally contribute to sinking of the system, they are comparatively lightweight organisms compared to the oysters.

## Conclusions

We have demonstrated that Jurassic crinoids did inhabit this unique pseudoplanktonic niche, becoming highly adapted, fast growing, very lightweight and self-sufficient viable communities. In the early Mesozoic seas, these rafts were home to far larger and more complex, now lost, ecosystems whose existence was a necessity as a result of a high number of anoxic shallow water basins (37, 38). Development of this lifestyle ensured the continued success of the group. The appearance of wood boring bivalves, and the prevalence of angiosperm wood along with the restoration of healthy benthic environments by the Middle Jurassic (9, 13) meant that this unique ecological phenomenon vanished from the fossil record forever.

## Methods

### Data collection

Photographs were taken from six specimens and composite images produced. These include small, medium and finally massive giant colonies from the following German museums and collections: Geoscience Centre of the University of Göttingen; Werkforum Museum, Dotternhausen; Naturmuseum Senckenberg in Frankfurt am Main; Geological Institute, University of Tuebingen; Staatliches Museum für Naturkunde, Stuttgart; and Urweltmuseum Hauff, Holzmaden.

### Spatial analysis

Randomly distributed points (Fig. 1*A*) can be modelled using homogeneous Poisson processes, and are found, for example, when larvae attach to an substrate not impacted by strong currents nor patchy environmental variables (39). While spatial processes that depend on position on the substrate can be modelled using heterogeneous Poisson processes, with density changing according to a given formula. To assess how disc density changed along the log, disc density was modelled as a heterogeneous Poisson process dependent on the *x* co-ordinate and then the *y*-co-ordinate. Model fit was assessed using the model residuals by plotting Q–Q and smoothed residual plots. If the observed line in the Q–Q plot fell outside two standard deviations of the model, the model was rejected (18, 40). Akaike information criterion values (41) were used to compare the relative quality of the statistical models that fitted the data.

### Moisture diffusion models

As it is not possible to know the actual properties of the fossil wood (due to the lack of fossil examples) or the actual growth rate of the oysters (as these species are now extinct), two types of models were developed. The first is a set of ‘catastrophic’ models (*SI Appendix*, Figs. S4– S9) which look at how long a series of size defined structures can survive with no natural sealing (see *Diffusion analyses* and *SI Appendix*). The second model is predictive, and proposes how long the mega-raft could have lasted if the system was efficiently sealed and a population of oysters (based in extant oysters) was growing on its surface (see *Oyster community growth model* and *SI Appendix*).

#### Diffusion analyses

For the first model, three sets of size defined colonies (*SI Appendix*, Figs. S4–S9)), including small colony specimens (S1 and S2) (*SI Appendix*, Figs. S4 and S5) medium colonies (M1 and M2) (*SI Appendix*, Figs. S6 and S7) and finally massive giant colonies from Holzmaden (G1 “The Hauff Specimen” and G2) (*SI Appendix*, Figs. S8 and S9) were tested. To address the stability of the log and colony, the density/diffusion model is constructed for each of the colony three-size classes using the log properties without any maximum growth population attached, then the crinoid community is added, along with a layer of oysters (11). The models use a colony weight based on modern isocrinid *Metacrinus* and Japanese oysters (*Crassostrea gigas*). They assume there is no natural sealing of the wood and it begins to absorb water, and therefore breakdown, from the onset of the colony (see *SI Appendix*).

#### Oyster community growth model

The second model considers the growth of the oyster community on the surface of the same logs and estimates how long the log can remain afloat while bearing the weight of the colony. It relies on oyster population data about extant communities of *Ostrea chilensis* and *Crassostrea virginica*. The model of population growth assumes the settlement on one single individual when the log contacts sea water and estimates the added mass of the community to the completely sealed log every 1-year increment until the whole log and colony sinks due to its density (see *SI Appendix*, Fig. S2 and Table S3).

## ACKNOWLEDGEMENTS

We thank the following institutions for access to the specimens: Geoscience Centre of the University of Göttingen; Werkforum Museum, Dotternhausen; Naturmuseum Senckenberg, Frankfurt am Main; Geological Institute, University of Tuebingen; Staatliches Museum für Naturkunde, Stuttgart; and Urweltmuseum Hauff, Holzmaden. We thank T Ewin, Natural History Museum, London, for his help with the early stages of the project. Also special thanks to R Haude, University of Göttingen, for providing the composite image of Holzmaden Log (G1) (Fig. 1*A* and *SI Appendix*, Fig. S1). We acknowledge support from DAAD funding enabling collection of data (A.W.H.) and a Junior Research Fellowship from Murray Edwards College, University of Cambridge (E.G.M.).

## Data availability

Extended methods for the spatial analysis and the diffusion analysis as well as properties, definitions and equations are discussed in detail in *SI Appendix*.

## Author contributions

A.W.H., D.C. and E.G.M. contributed equally to research, discussion, and manuscript preparation. A.W.H designed the research project, collected the museum data, photographed fossil material and prepared the specimen figures including the spatial analysis plots. D.C. designed and ran the diffusion analyses, and prepared the corresponding diagrams. E.G.M. designed and ran the spatial analysis, and prepared the corresponding diagrams. C.M. prepared the reconstruction.

The authors declare no conflict of interest.

