## Supplementary Information for "Reconstructing the ecology of a Jurassic pseudoplanktonic megaraft colony"

**Supplementary Information: Reconstructing the ecology of a Jurassic psedoplanktonic megaraft colony**

### Introduction to Crinoid Ecology

The crinoids that are the focus of this study are a wonder of the Mesozoic with as many as 100 individuals covering oyster-encrusted logs up to 14m long (*SI Appendix*, Fig. S1). The crinoids that inhabited these communities belong to distinct genera with a characteristic morphology. Although a number of unique adaptations have been suggested for these animals, uncertainty exists whether or not this mode of life was possible. Our study is consistent with the special adaptations that these crinoids have, such as distally tapering column, a strengthened attachment structure and the development of an enlarged crown, which does not need to close hermetically in *Seirocrinus*, and a very large crown and high densities of cirri allowing for the enlargement of the cup and the food groove in *Pentacrinites*. Never again do crinoids develop such adaptations to an unusual ecosystem. Although risky, this adaptation allowed these crinoids to spread across Eurasia inhabiting regions as widespread as Alaska and Japan (Hunter and Zonneveld 2008, Hunter et al. 2011).

### 1. Analysis of the spatial positions of attachment discs along the Holzmaden (G1) crinoid log

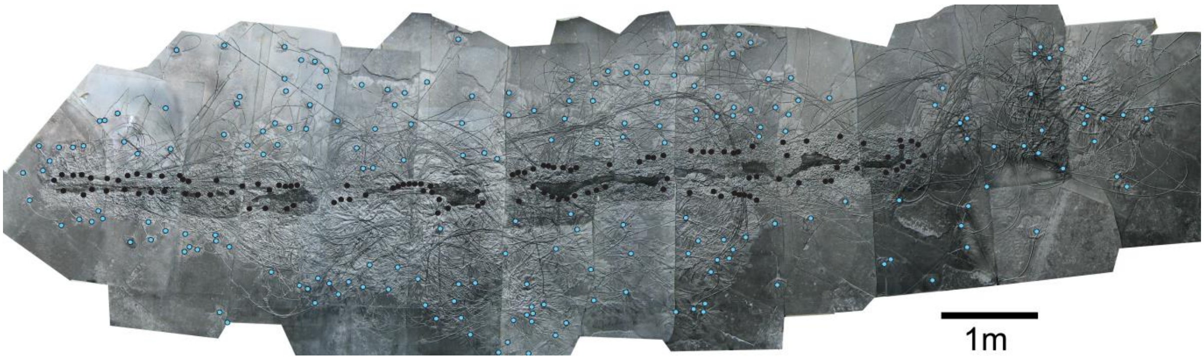


**Fig. S1.** The specimen with attachment discs marked in black and heads marked in blue.

#### 1. 1 Introduction to SPPA

This species of crinoid are immobile once attached to logs or other substrate (Hagdorn 2016). Therefore, their spatial distributions of their attachment discs on their settled substrate reflect the dispersal processes that brought their larvae to the substrate and the consequent interactions between organisms. As such, analysis of the spatial distributions using spatial point process analyses (SPPA) can shed light on these dispersal process and interactions (e.g. Seidler and Plotkin 2006). SPPA has been effectively applied to the study of modern sessile ecosystems, most notably in terrestrial forests, but also to a limited extent with marine benthic organisms such as barnacles (Lancaster and Downes 2004, Illian et al. 2008).

Comparisons with extant organisms such as barnacles show that when larvae attach to a moving object, such as a boat hull, the spatial distributions are highly anisotrophic - that is they are not uniform across the entire object. Areas which face into the current (such as the bow of a boat) are subject to high turbulence and thus have limited colonization, while the areas sheltered from the current, such as boat sterns, have the highest densities of invertebrate colonizers (Ashton et al. 2014). In contrast when larvae attach to an immobile substrate, their spatial distributions exhibit complete spatial randomness (CSR) when not impacted by environmental variables such as habitat heterogeneities such as patchy rocks (e.g. Edwards and Stachowicz 2011). Comparisons of observed spatial distributions with spatial models can be used to infer the most likely process underlying a spatial distribution. Randomly distributed points, (CSR) can be modelled using homogeneous Poisson processes while anisotrophy can be modelled using heterogeneous Poisson processes. For these heterogeneous Poisson processes, the density of points changes according to a formula, such as with distance along the substrate.

Pair correlation functions (PCFs) are commonly used to describe complex spatial distributions over large distances, where they document how the density of specimens changes with distance (e.g. Wiegand et al. 2007). A CSR (random) population will have a PCF of 1, whereas aggregation is indicated by PCF >1, and segregation by PCF <1. The magnitude of the PCF reflects the intensity of biological and physical processes; a population with PCF=4, for example, is four times more aggregated than one with CSR. If a PCF is significantly non-random, then the specimens have been subject to a biological or ecological process, such as interactions with each other, or their environment. Segregations between specimens occur when organisms cannot overlap with each other, or being within the vicinity of another specimen has a negative effect, while aggregations indicate likely positive effects, such as beneficial substrate, or dispersal induced clustering.

#### 1.2 Methods

Data exploration, inhomogeneous Poisson modelling and residual analysis was performed in R using the package spatstat (R core team, 2013, Baddeley et al. 2000, Baddeley et al. 2015). Pair correlation functions (PCFs) were calculated to describe the spatial distributions of discs on the log (*SI Appendix*, Fig. S1) (Illian et al. 2008). Monte Carlo and Diggle's goodness-of-fit test5 (the p-value $p_d$, in which $p_d$ =1 indicates a perfect model fit, and $p_d$ =0 indicates no fit), simulations were used to assess whether the spatial distribution was completely spatially random (Diggle 2003). The PCF value reflects how many times more likely the distribution seen is aggregated (or segregated) compared with CSR. The PCF was plotted (*SI Appendix*, Fig. S2), and nine hundred and ninety-nine simulations were run to generate simulation envelopes around the CSR. To assess whether the density of the spatial distributions of discs was stronger in any particular direction (that is, it exhibited isotropy); density plots were fit to of the point positions of the discs. To assess how disc density changed along the log, disc density was modelled as a heterogeneous Poisson process dependent on the *x* co-ordinate and then the *y*-co-ordinate. Model fit was assessed using the model residuals (Illian et al. 2008, Wiegand et al. 2007). Model residuals assessed the fit of the model to the data by plotting Q–Q and smoothed residual plots. If the observed line in the Q–Q plot fell outside two standard deviations of the model, the model was rejected (Illian et al. 2008, Wiegand et al. 2007).

Akaike information criterion values (Baddeley et al. 2011) were used to compare the relative quality of the statistical models that fitted the data.

#### 1.3 Results

The spatial distribution of the discs was found to be significantly segregated ($p_d$ <0.001) below 4.5cm (*SI Appendix*, Fig. S2). This segregation is hard-core under 2.5cm, which means that no specimens are found within 2.5cm of each other. Between 2.5 and 4.5cm there is a soft-core segregation, which means that while segregation occurs, the likelihood of specimens occurring within this distance is reduced. This result suggests that the attachment discs require a non-overlapping area of 2.5cm, and it is sub-optimal to attach within 2cm of another disc. The density map shows a clear anisotrophy: that is a difference in density dependent on direction, with the highest density on the left hand side, with decreasing density along the log (*SI Appendix*, Fig. S3). This anisotrophy can be modelled by heterogeneous Poisson model depending on the *x* co-ordinate (*SI Appendix*, Table S1).

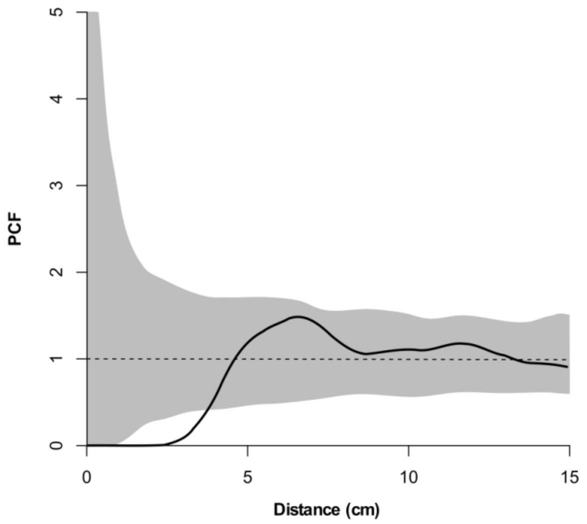


**Fig. S2.** Pair correlation function for attachment discs. The x axis is the inter-point distance between organisms in centimetres. On the y axis, PCF=1 indicates CSR, <1 indicates segregation and >1 indicates aggregation. Grey shaded area depicts the bounds of 99 Monte Carlo simulations of CSR. Since the PCF curve is not completely within these areas, the CSR hypothesis is rejected and one can assume that discs are significantly segregated.

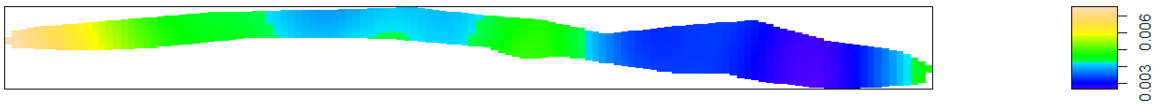


**Fig. S3.** Density map of attachment discs. Note the higher densities on the left hand side.

**Table S1.** AIC values for different heterogeneous Poisson models dependent on either the x or y co-ordinate. The lower the AIC value, the better fit the model. The lowest AIC value was for the heterogeneous Poisson model dependent on the x coordinate. This best-fit model was within the two standard deviations for the Q-Q plot.

| **Model** | **AIC** |
|---|---|
| CSR | 1592.544 |
| *X* | 1591.102 |
| *Y* | 1594.313 |

**2. Diffusion model for the Holzmaden crinoid log (*SI Appendix*, Fig. S1)**

We construct the model around a cylinder with longitudinal diffusion.

Using Fick's Law of diffusion =

$$C(t+1,i) = C(t,i) + \Delta_t \times D_{x} \left[ \frac{C_{i+1} - 2C_i + C_{i-1}}{\Delta^2} \right]$$

The C means water concentration; delta t is the duration between each time step (one day in our model); Dx is the diffusion coefficient in the longitudinal direction (which is a function of the conduit diameter and the moisture content); Delta is the distance increment between two slice compartments of the log). The growth of oysters adds to the mass and the population growth is modeled using a logistic curve.

### 2.1 Physical properties of wood

#### 2.1.1 Wood types and variety

Wood is an anisotropic material composed of dead and living cells through which moisture is transported. Due to the hygroscopic nature of this material the physical properties of wood, including its moisture transport properties, are strongly influenced by both the surrounding moisture as well as the ambient temperature (Skaar 1984). Three directions are considered when examining tree wood: the longitudinal, radial and tangential directions. The longitudinal direction is the direction of the wood fibers. The radial direction runs from the center of the tree to the outside. Finally, the tangential direction follows the curvature of the tree (Jacobson and Banerjee 2006). Trees are divided into two classes: conifers (or softwoods) and broadleaf trees (or hardwoods). The main distinction between the two classes is that hardwoods possess vessel elements in addition to the common wood fibers (or tracheids). Both classes display a broad range of densities with, for example, balsa (hardwood) showing lower densities ($0.2e10^3$ $kg/m^3$) than softwoods and yew (softwood) being denser ($0.7e10^3$ $kg/m^3$) than many hardwoods.

#### 2.1.2 Structures

Cross sections for hardwoods and softwoods reveal dark and light areas that correspond to latewood and earlywood respectively. Each area is composed of tubular structures called wood fibers or tracheids that are around 30 μm in diameter and 3 mm in length (Jacobson 2006). The main structural distinction between softwood and hardwood is the existence of additional specialized vessels in the latter through which moisture is conducted. These vessels present a variety of features (presence or absence of perforations, size of the perforations, lignin reinforcements, etc.) and diameters. There is currently no definitive model that would consider the chemical and physical processes involved in moisture transport in wood and the phenomenon is likely to be affected by many variables simultaneously (Skaar 1984; Wood et al. 2002).

#### 2.1.3 Composition

Wood consists of three polymers: cellulose, hemicellulose and lignin. These polymers are not evenly distributed as cellulose amounts to between 40 and 50% of the material in mass, depending on the species (Sjostrom 1993). The remaining mass is roughly evenly divided between hemicelluloses and lignin. The three polymers assemble to form microfibrils that are typically 5000 nm long and 10-20 nm wide. In turn, these microfibrils are arranged in sheets that constitute the cell wall of wood fibers and the relative thickness of these sheets is also species-specific. The varying abundance of the polymers coupled with their distinct chemical properties explain the complex sorption behaviour of wood materials.

#### 2.1.4 Wood Decay

It is unknown whether the log structures in this study were gymnosperms or angiosperms as the wood is not preserved; additionally, the variety of gymnosperm species themselves are critical to the long term survival of the raft. Generally recent gymnosperm confer forests produce much more wood debris than angiosperm forests, therefore we can assume there was plenty of available material. However, the structure of the wood itself and the agents of decay in the environment the wood is found are critical for the longevity of the system and how the log structure will break up. Firstly, the flow of the water will break the bark which acts as a protective coating around the wood increasing the infiltration rate; this will be exacerbated by natural breaks or cracks; alternatively the wood could have been crushed. However, the log structures clearly survived high-energy fluviatile environment or entered the system from the margins of the epicontinental sea. From this, we can infer that a log from the Jurassic gymnosperm forest in a swamp, estuary or delta was dislodged and floated out to sea where oysters, other sealing agents and crinoid larval disks attached and formed a complex community structure. Fresh water fluviatile environments have surprisingly lower rates of microbial decay in streams compared to the open ocean; conversely these environments are home to elevated amounts of fungus and terrestrial invertebrates such as insects. The latter could not survive in the open ocean environment finding it intolerable. It is often cited that log structures are less common in the open ocean due the presence of marine invertebrates that would break down the wood structures; however, our examples were likely protected by the oysters and evidence of anoxia prevalent in both the Holzmaden and in the Lias may have meant few invertebrates could have infiltrated the log. Additionally, many of the agents found in the modern oceans that break down floating wood evolved post mid-Jurassic. In terms of properties of the wood itself, the initial density can influence decay rates. Our idealized models assume that the density of wood was the same across the log; close to the composition of sapwood that contains functioning vascular tissues. Therefore spaces are likely to absorb water and decay at a higher rate. Gymnosperm tracheids (pore spaces) are much smaller than angiosperms vessels. The density of these spaces in gymnosperm wood declines higher up the tree structure which means the low density structures further up the log would have been better candidates for longer lasting wood megarafts. This sapwood is surrounded by outer bark, inner bark which includes the phloem and core consisting of the heartwood. With a much lower rate of absorption the presence of plentiful heartwood would have strengthened the viability of the system considerably. Gymnosperm sapwood contains much less living tissue 5-11 % as opposed to 11-48 % found in angiosperms. The lack of living tissue would have meant a much lower decay rate in this type of wood.

### 2.2 Moisture sorption model

#### 2.2.1 Basis of the model

A Fickiandiffusion model was used to evaluate the soaking time of wood logs (Jacobson and Banerjee 2006, Wood and Gladden 2002). Debate is still open concerning the Fickian nature of the moisture transport as water appears in three distinct phases in wood: free, bound and gaseous (Krabbenhøft et al. 2004). However, as there is currently no model suited to the Non-Fickian behaviour in wood (Wadsö 1994), we decided to use a Fickian model to provide an order of magnitude for the soaking durations until sinking. Despite being extremely simplified, it has been shown to be equivalent to other models at least in isothermal conditions (Krabbenhøft et al. 2004). Previous research has shown that the Fickian model systematically underestimates the value of diffusion coefficients mostly during the initial phase of absorption (Shi 2007), which means that it predicts soaking speeds that are actually slower than the experimental measurements. Therefore, the estimate times to sinking considered in this study should be considered as maximum values.

#### 2.2.2 Diffusion model

This model assumes that the wood is green with a moisture content higher than the fiber saturation point so as to avoid any impact of volume change for the vessels. As the free water is already in the vessels, most of the soaking is the result of the diffusion of either water bound to the cell walls (bound water) or vapor. The present two-phase model examines the diffusion of moisture under isothermal conditions using a finite difference technique to solve the differential equations that involve diffusion coefficients that are varying according to the moisture content. The model is considering an isothermal diffusion because heat diffusivity is orders of magnitude higher than moisture diffusivity in wood.

#### 2.2.3 Diffusion coefficients

The diffusion of gas occurs within the lumen of cells and is dependent on the diffusivity of water vapor in air, temperature and the saturated vapor pressure. The diffusion of bound water is dependent on the temperature and the moisture content. These two values can be combined to reflect the general diffusion coefficient (Baronas et al. 2001).

#### 2.2.4 Moisture movement in wood

The model considers the diffusion of moisture within a cylinder of revolution of radius *r* and length *L*. Wood vessels are running along the length of the cylinder and are orthogonal to the circular cross section.

The diffusion of moisture can be modeled as a Fickian phenomenon by the equation (1) that is set in a one-dimension (longitudinal) system without convection.

$$\frac{\partial u}{\partial t} = D\frac{\partial^2 u}{\partial x^2} \quad (1)$$

In a cylindrical system, the equation should be adapted to reflect the diffusion that happens both longitudinally and radially:

$$\frac{\partial u}{\partial t} = D_r\left[\frac{1}{r}\frac{\partial}{\partial r}\left(r\frac{\partial u}{\partial r}\right)\right] + D_l\frac{\partial^2 u}{\partial x^2} \quad (2)$$

Initial conditions

u(0,r,t)=u(L,r,t)=100; u(x,b,t)=100;

#### 2.2.5 Longitudinal component

If we employ a classic separation method, we can express $u_l$(r,x) as a product of two functions X and T such as:

$$u_l(r,x) = X(x)T(t) \quad (3)$$

The longitudinal component of equation (2) becomes:

$$\frac{1}{D_l}\frac{T'}{T} = \frac{X''}{X} = -\mu \quad (4)$$

If we consider the spatial component of this equivalence first, we can say that μ>0 is the only condition leading to non-trivial solutions:

| | | |
|---|---|---|
| | $X_n(x) = A_n \sin\left(\frac{n\pi x}{L}\right)$ | (5) |

with $\mu=(n\pi/L)^2$ . By injecting this value into the temporal component, we obtain:

| | | |
|---|---|---|
| | $T_n' + \frac{D_l n^2 \pi^2}{L^2} T_n = 0$ | (6) |

which leads to:

| | | |
|---|---|---|
| | $T_n(t) = B_n \exp\left(\frac{-D_l n^2 \pi^2}{L^2} t\right)$ | (7) |

By combining (5) and (7) and summing all the components, we obtain a general expression for the moisture profile along the longitudinal axis:

| | | |
|---|---|---|
| | $u_l(x,t) = 100 - \sum_{n=1}^{\infty} C_n \sin\left(\frac{n\pi x}{L}\right) \exp\left(\frac{-D_l n^2 \pi^2}{L^2} t\right)$ | (8) |

the value of $C_n$ is fixed by the initial condition at t=0 and the boundary conditions at x= L:

| | | |
|---|---|---|
| | $C_n = \frac{2}{L} \int_0^L \varphi(x) \sin\left(\frac{n\pi x}{L}\right) dx$ | (9) |

For example, if $L=\pi$ and $u_l(x,0)=-x(\pi-x)$,

| | | |
|---|---|---|
| | $C_n = \frac{2}{L} \int_0^{\pi} x(\pi - x) \sin(nx)\, dx$ | (10) |
| and | $C_n = 4 \frac{1-(-1)^n}{n^3 \pi}$ | (11) |

This initial distribution of the moisture is a good approximation of a moisture profile.

In the end, the longitudinal diffusion of moisture in a cylinder of length $\pi$ can be expressed as:

| | | |
|---|---|---|
| | $u_l(x,t) = 100 - \frac{8}{\pi}\sum_{n=1}^{\infty}\frac{\sin\big((2n-1)x\big)}{(2n-1)^3}\exp(-D_l(2n-1)^2 t)$ | (12) |

#### 2.2.6 Radial component

A similar approach can be taken with the radial component of the moisture function.

| | | |
|---|---|---|
| | $\frac{\partial u_r}{\partial t} = D_r\left(\frac{\partial^2 u}{\partial u^2} + \frac{1}{r}\frac{\partial u}{\partial r}\right)$ | (13) |

A separation of the variables leads to:

| | | |
|---|---|---|
| | $u_r(r,t) = R(r)T(t)$ | (14) |

and thus equation (12) can be re-written as:

| | | |
|---|---|---|
| | $\frac{1}{D_r}\frac{1}{T}\frac{\partial T}{\partial t} = \frac{1}{R}\left(\frac{\partial^2 R}{\partial r^2} + \frac{1}{r}\frac{\partial R}{\partial r}\right) = -\left(\frac{m}{b}\right)^2$ | (15) |

The only non-trivial solutions for equation (14) are for a separation constant that is negative ($-(m/b)^2$) and this leads to:

| | | |
|---|---|---|
| | $R(r) = J_0\left(\frac{m}{b}r\right)$ | (16) |

If we focus now on the temporal element of the moisture function, the boundary condition R(b)=0 suggests to consider all the possible solutions of the equation $J_0(m_n)=0$

The equation (14) becomes then:

| | | |
|---|---|---|
| | $\frac{\partial T_n}{\partial t} + \left(\left(\frac{m_n}{b}\right)^2 D_r\right)T_i = 0$ | (17) |
| and thus | $T_n(t) = M_n \exp\left(-\left(\frac{m_n}{b}\right)^2 D_r t\right)$ | (18) |

By combining (15) and (17) and summing all the components, we obtain a general expression for the moisture profile along the longitudinal axis:

| | | |
|---|---|---|
| | $$u_r(r,t) = \sum_{n=1}^{\infty} M_n J_0\left(\frac{m_n}{b} r\right) \exp\left(-\left(\frac{m_n}{b}\right)^2 D_r t\right)$$ | (19) |

The boundary condition $u_r(r,0)=M_{init}$ leads to:

| | | |
|---|---|---|
| | $$M_n = \frac{(2M_{init})}{m_n J_1(m_n)}$$ | (20) |

and therefore:

| | | |
|---|---|---|
| | $$u_r(r,t) = 2M_{init} \sum_{n=1}^{\infty} \frac{1}{m_n J_1(m_n)} J_0\left(\frac{m_n}{b} r\right) \exp\left(-\left(\frac{m_n}{b}\right)^2 D_r t\right)$$ | (21) |

**2.2.7 Weight of the Colony**

Part of our calculation is to seek to understand the total weight of the colony. The flowing factors need to be taken into account: Firstly, the maximum carrying capacity of the log (see *SI Appendix*, oyster community growth model below) and secondly the original density of the wood material. We take into account the following factors:

1) The population is always preserving optimum intrinsic growth parameters.

2) No environmental factor limits the growth of the community during the duration of the simulation.

3) There is no significant water absorption within the oysters or the crinoids during the growth of the colony.

We conclude that the mass of the community in the most extreme circumstances could reach around 880 kg for a log that is 10 m long and 0.4 m in diameter with a carrying capacity of 1000 individuals per square meter. If the carrying capacity is less than 600 individuals per square meter, only logs with a density higher than 0.5 (pine is around 0.5) would manage to sink within a 20-year window and the community would then weigh around 630 kg. As for the mass of the log a Shortleaf pine (softwood) with a kiln dry specific gravity of 0.47 (average for softwood), the green wood will have a general mass varying between 950 and 1100 kg for a log that is 10 m long and 40 cm in diameter. This could be regarded as a minimum value with modern hardwood logs up to 15,000 kg. That means that the weight of the oyster community in that optimum growth scenario (super high carrying capacity) matches almost the original mass of a softwood log. However these high carrying capacities >200 hundred oysters per square meter are rare and it is likely that the oyster community mass did not reach 600 kg for a span of 20-50 years on our large logs.

### 2.3 Results

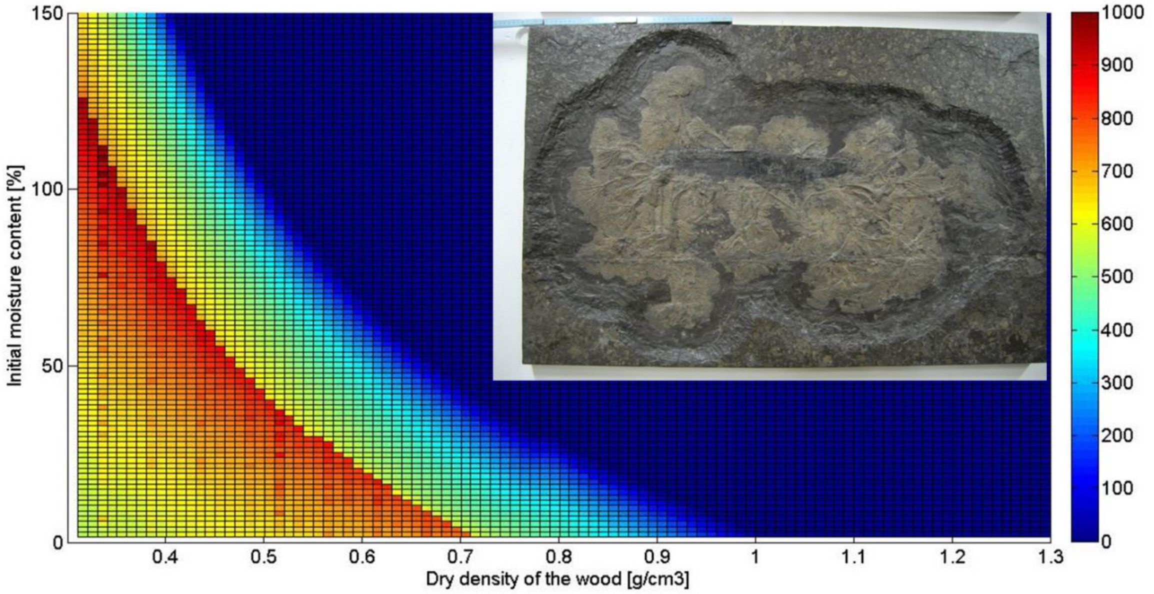


**Fig. S4.** Small colonies - Göttingen (S1) and Stuttgart 1 (S2) – 40 cm long Diameter 5-6 cm (Community Removed).

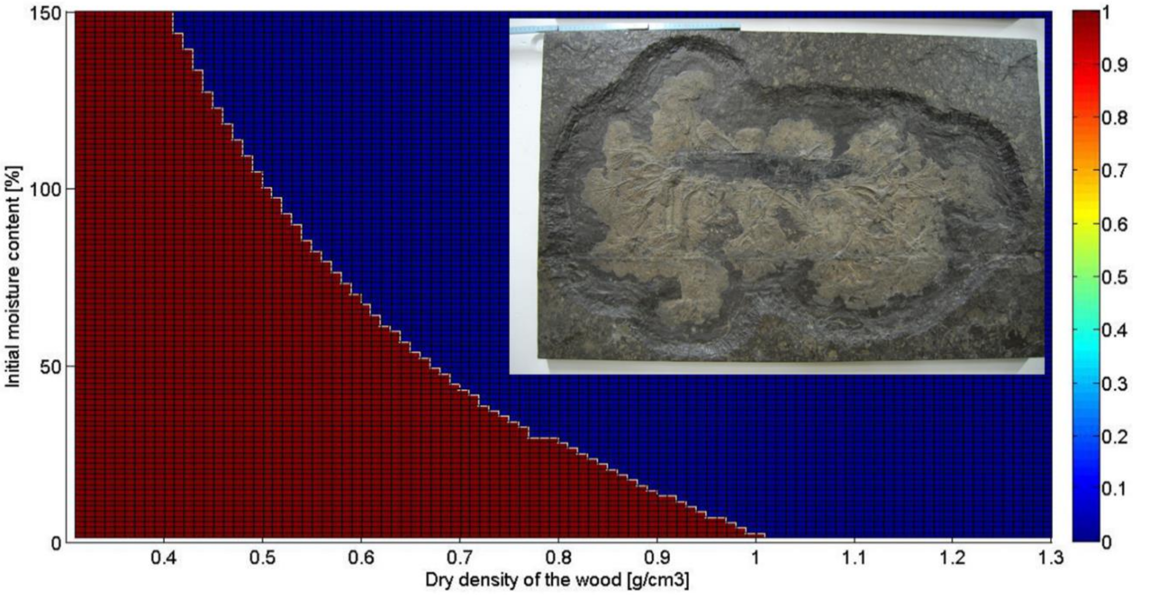


**Fig. S5.** Small colonies - Göttingen (S1) and Stuttgart 1 (S2) – 40 cm long Diameter 5-6 cm (Community Included).

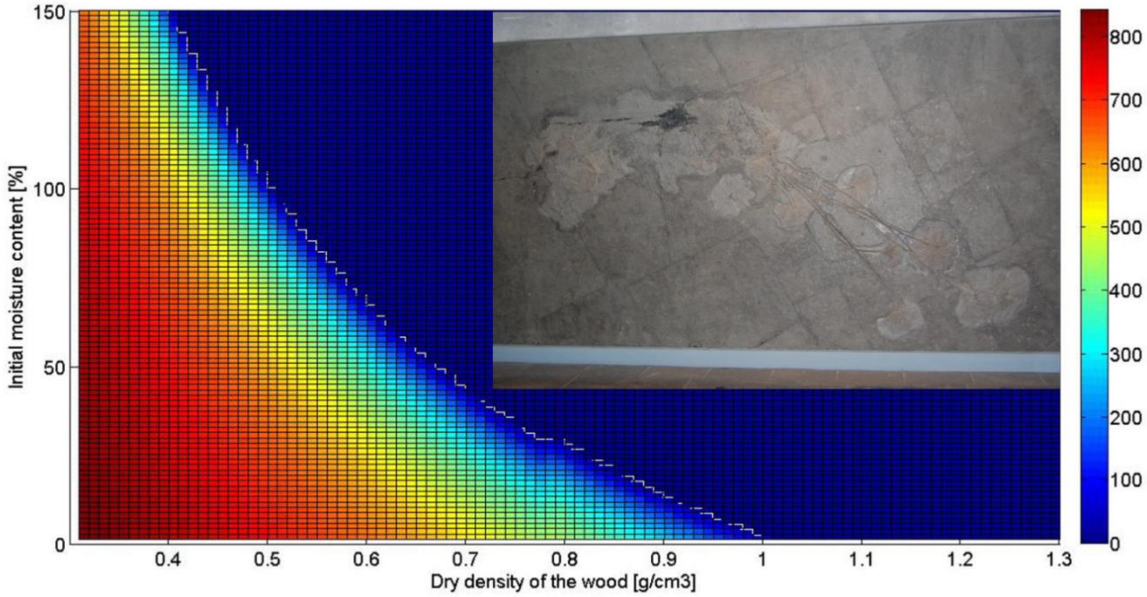


**Fig. S6.** Medium colonies - Frankfurt (M1) and Dotternhausen (M2) – 190 cm long, Diameter 8-12 cm (Community Removed).

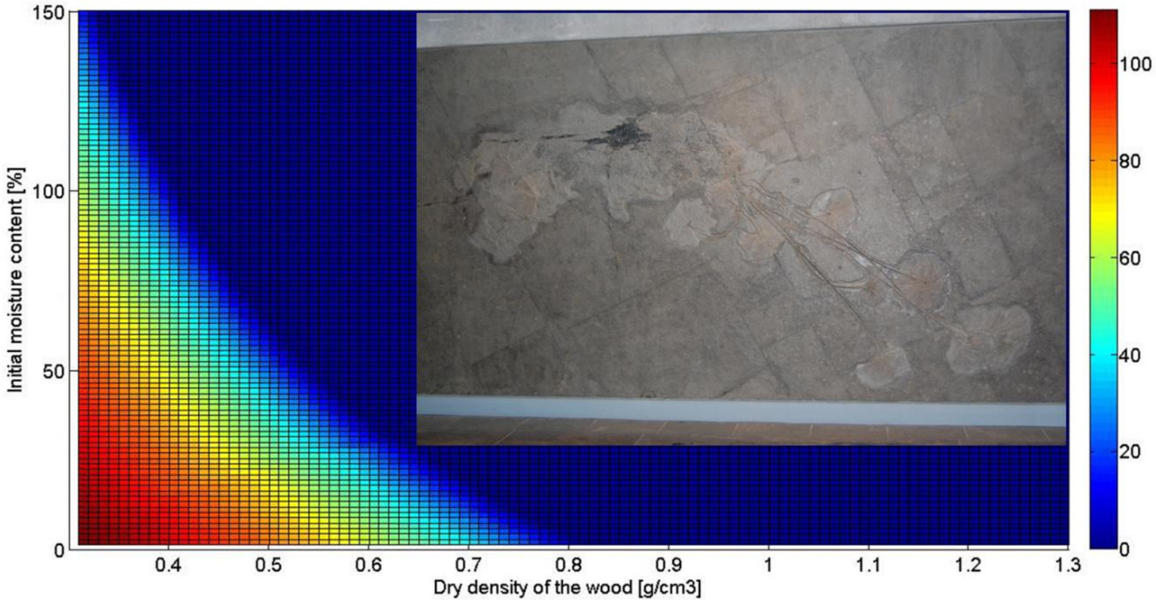


**Fig. S7.** Medium colonies - Frankfurt (M1) and Dotternhausen (M2) – 190 cm long, Diameter 8-12 cm (Community Included).

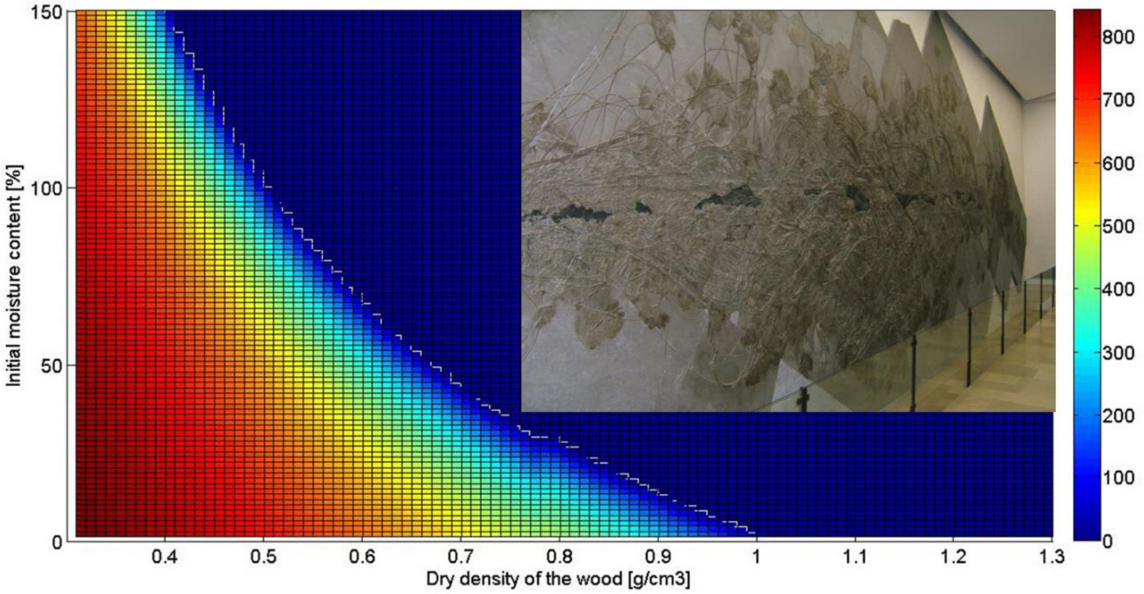


**Fig. S8.** Massive colonies – Holzmaden (G1) and Stuttgart 2 (G2) – 12 m long Diameter 25-28 cm (Community Removed).

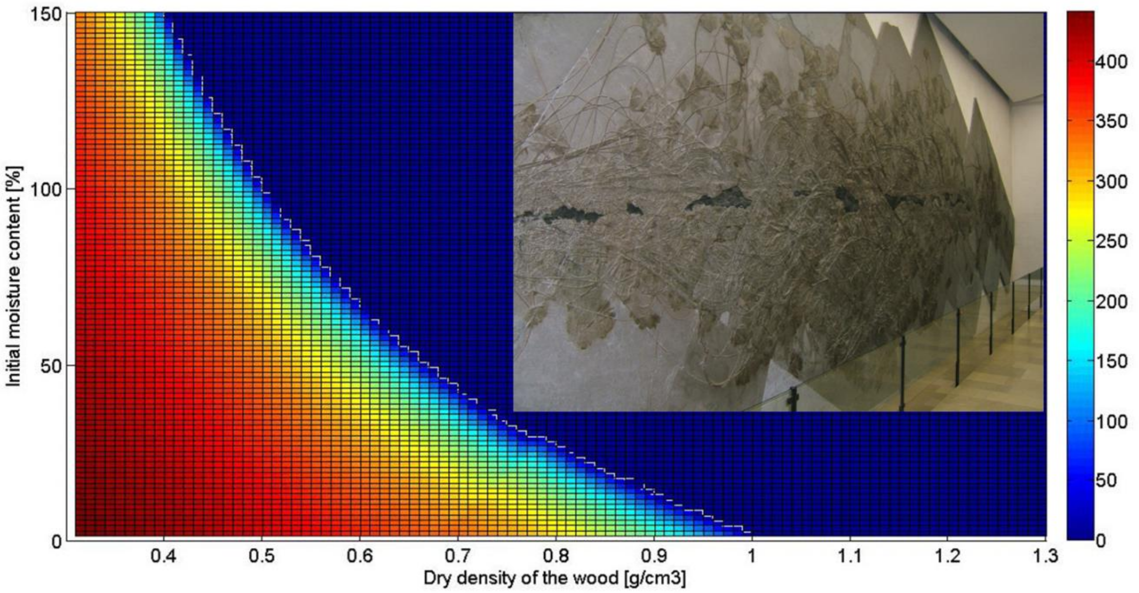


**Fig. S9.** Massive colonies – Holzmaden (G1) and Stuttgart 2 (G2) – 12 m long Diameter 25-28 cm (Community Included).

### 3 Oyster community growth model

#### 3.1 Introduction

The population model incorporated both the spatial distribution along the log as well as life-history estimates (fecundity, mortality, settling rates and maturation time) for a complete life cycle of extant analog *Ostrea chilensis* (Philippi, 1845). A life cycle was defined as the duration from the release of eggs, through the planktonic larval stage, settlement and growth to reach a mature stage allowing the release of the next generation of eggs. Simulations for the growth of oyster communities were conducted using the mathematical package MATLAB R2017a. The log surface was sectioned into 10 compartments (or boxes) and the model quantitatively estimated the success (settlement and growth) of a population within a given box (Kanary et al. 2011). The model was initialized by assuming one mature adult in box 1 and it estimated the number of mature oysters at each following time increment (one year) across the log (10 boxes).

#### 3.1 Oyster Diffusion Model

The model assumes that once settled, an oyster does not relocated between boxes (Raillard and Ménesguen 1994) and that the time to reach maturity is 1 year (Lannan 1971). In a population, only a certain percentage is actually spawning (rSpawn) which leads these mature adults to release eggs (per capita reproductive rate rRep). Of these released eggs, only a fraction of the larvae can settle (sRate). The further the box is away from the spawning location, the less likely the larva will settle and a simple transition matrix $tP_{(I,n)}$ was designed to illustrate this differential implantation distribution. Each entry in the transition matrix $tP_{(I,n)}$ represents the probability of transitioning from one compartment to another through simple dispersal by water currents because the average swimming speed of a veliger larva only ranges between 0.6 and 2 cm/min (Hidu and Haskin 1978). These assumptions provided a simplified framework for the growth model.

The population growth process can thus be expressed as:

$$\boldsymbol{X_{(i,t)}} = (\boldsymbol{1} - \boldsymbol{r_{Mort}})\boldsymbol{X_{(i,t-1)}} + \sum_{n=1}^{10} \boldsymbol{r_{Spawn}}\,\boldsymbol{r_{Rep}}\boldsymbol{s_{Rate}}\left(\boldsymbol{tP_{(i,n)}} * \boldsymbol{X_{(n,t-1)}}\right) \qquad \textbf{(1)}$$

With

$$\boldsymbol{X_{i,t}} = \boldsymbol{min}\{\boldsymbol{X_{i,t}}, \boldsymbol{K}\} \qquad \textbf{(2)}$$

Where K is the maximum carrying capacity of the log compartment (individuals/ $m^2$). Carrying capacity is understood as a simple saturation function: at each time step, if the value of X computed by equation 1 is larger than K, it is truncated to K because additional larvae cannot settle on a log compartment once the carrying capacity has been reached. In addition, n is the index associated to the destination compartment.

Spawning rate, per capita reproductive rate and mortality rate correspond to average values taken from the literature about *Ostrea chilensis* (Brown 2011) or *Crassostrea virginica* (Powell et al. 1996, Wang et al. 2008). The values for the parameters appear in *SI Appendix*, Table S2.

| Parameter | Value | Reference |
|---|---|---|
| **Mortality rate (rMort)** | **0.6** | **(Brown 2011)** |
| **Spawning rate (rSpawn)** | **0.15** | **(Wang et al. 2008)** |
| **Per capita reproductive rate (rRep)** | **500** | **(Brown 2011)** |
| **Settlement rate (sRate)** | **0.10** | **(Powell et al. 1996)** |
| **Maximum carrying capacity (K)** | **50 ~ 1000 individuals / $m^2$** | **(Brown 2011)** |

**Table S2.** Values of the parameters retained for the model of population growth. Values are extracted from existing literature about oyster communities in New Zealand (Brown 2011) and the USA (Powell et al. 1996, Wang et al. 2008).

The model allowed to reconstruct the number of years required to reach sinking density for the whole log for various values of K (*SI Appendix*, Table S3). In some cases, the dimensions of the log combined with the density of the wood material would allow the loaded log never to reach sinking threshold within the 20-year window.

| Wood density | | Maximum carrying capacity (population density in ind/m2) | | | | | | | | | | | | | | | | | | | |
|---|---|---|---|---|---|---|---|---|---|---|---|---|---|---|---|---|---|---|---|---|---|
| | | 50 | 100 | 150 | 200 | 250 | 300 | 350 | 400 | 450 | 500 | 550 | 600 | 650 | 700 | 750 | 800 | 850 | 900 | 950 | 1000 |
| | 0.3 | x | x | x | x | x | x | x | x | x | x | x | x | x | x | x | x | x | 6 | 6 | 6 |
| | 0.4 | x | x | x | x | x | x | x | x | x | x | x | x | x | x | x | 6 | 6 | 6 | 6 | 6 |
| | 0.5 | x | x | x | x | x | x | x | x | x | x | x | x | 6 | 6 | 6 | 6 | 6 | 6 | 6 | 6 |
| | 0.6 | x | x | x | x | x | x | x | x | x | x | 5 | 5 | 5 | 5 | 5 | 5 | 5 | 5 | 5 | 5 |
| | 0.7 | x | x | x | x | x | x | x | 5 | 5 | 5 | 5 | 5 | 5 | 5 | 5 | 5 | 5 | 5 | 5 | 5 |
| | 0.8 | x | x | x | x | x | 5 | 5 | 5 | 5 | 5 | 5 | 5 | 5 | 5 | 5 | 5 | 5 | 5 | 5 | 5 |
| | 0.9 | x | x | 5 | 5 | 5 | 5 | 5 | 5 | 5 | 5 | 5 | 5 | 5 | 5 | 5 | 5 | 5 | 5 | 5 | 5 |
| | 1.0 | 1 | 1 | 1 | 1 | 1 | 1 | 1 | 1 | 1 | 1 | 1 | 1 | 1 | 1 | 1 | 1 | 1 | 1 | 1 | 1 |

**Table S3.** Number of years required for a growing population of oysters to sink a sealed wood log. In this simulation, the wood log had a constant geometry with a length of 10 m and a radius of 0.2 m, corresponding roughly to the dimensions of the Holtzmaden log.
